# PIGNON: A protein-protein interaction-guided functional enrichment analysis for quantitative proteomics

**DOI:** 10.1101/2021.02.03.429592

**Authors:** Rachel Nadeau, Anastasiia Byvsheva, Mathieu Lavallée-Adam

## Abstract

**Background:** Quantitative proteomics studies are often used to detect proteins that are differentially expressed across different experimental conditions. Functional enrichment analyses are then typically used to detect annotations, such as biological processes that are significantly enriched among such differentially expressed proteins to provide insights into the molecular impacts of the studied conditions. While common, this analytical pipeline heavily relies on arbitrary thresholds of significance. Indeed, a functional annotation may be dysregulated in a given experimental condition, while none or very few of its proteins may be individually considered to be significantly differentially expressed. Such an annotation would therefore be missed by standard approaches.

**Results:** Herein, we propose a novel graph theory-based method, PIGNON, for the detection of differentially expressed functional annotations in different conditions. PIGNON does not assess the statistical significance of individual genes, but rather maps protein differential expression levels onto a protein-protein interaction network and measures the clustering of proteins from a given functional annotation within the network. This process allows the detection of functional annotations for which the proteins are differentially expressed and grouped in the network. A Monte-Carlo sampling approach is used to assess the clustering of proteins in an expression-weighted network. When applied to a quantitative proteomics analysis of different molecular subtypes of breast cancer, PIGNON detects Gene Ontology terms that are both significantly clustered in a protein-protein interaction network and differentially expressed across two breast cancer subtypes. PIGNON identified 168 breast cancer pathways dysregulated and clustered within the network between the HER2+ and triple negative subtypes, 203 breast cancer pathways shared by HER2+ and hormone receptor positive subtypes, 19 breast cancer pathways shared by hormone receptor positive and triple negative breast that are not detected by standard approaches. PIGNON identifies functional annotations that have been previously associated with specific breast cancer subtypes as well as novel annotations that may be implicated in the diseases.

**Conclusion:** PIGNON provides an alternative to functional enrichment analyses and a more comprehensive characterization of quantitative datasets. Hence, it contributes to yielding a better understanding of dysregulated functions and processes in biological samples under different conditions.

## BACKGROUND

High throughput quantitative proteomics studies provide an overview of the activity of different processes and functions in a biological sample. Mass spectrometry-based quantitative proteomics approaches are routinely used to quantify proteins in biological samples[1–4]. Strategies involving both stable isotope labelling[5–8] and label-free mass spectrometry[9] can be used to compare the expression of proteins across different samples and experimental conditions. When contrasting protein expression in different experimental conditions, it is common to use univariate statistical tests, such as the Student’s t-test and Mann Whitney U test to assess the significance of protein differential expression. Arbitrary thresholds are then typically established (e.g. a p-value < 0.05, expression fold-change > 1.5 or < 0.66) to determine the set of proteins deemed differentially expressed. Functional enrichment analyses are then often applied to the set of differentially expressed proteins in order to provide insights into the role they play in the biological samples and about the molecular impacts of the studied experimental conditions. Functional annotations investigated may take the shape of Gene Ontology (GO) terms[10], KEGG[11] and REACTOME[12] pathways or MSigDB signatures[13].

Computational tools such as Ontologizer[14], GoMiner[15], and DAVID[16] are often applied to compare the occurrences of the functional annotations among the set of differentially expressed proteins against all those in the entire set of quantified proteins using statistical tests such as the Fisher exact test to determine enrichment significance of the enrichment. Such a strategy is limited by the use of arbitrary cut-offs determining whether a protein is significantly differentially expressed or not. Proteins may achieve very similar significance values, but if each sits on different sides of the established cut-offs, they end up being treated very differently. Furthermore, many proteins sharing a given annotation may show some level of differential expression but not sufficient pass the chosen threshold. Such an annotation would therefore not be considered by the functional enrichment analysis even though a moderate differential expression of many proteins in a given pathway could contribute to the dysregulated phenotype. Methods such as Gene Set Enrichment Analysis (GSEA)[13] and GOrilla[17] have been proposed to counter such a loss of information by assessing whether genes sharing a given annotation are enriched towards the top or bottom of a ranked list based on differential expression. Similarly, PSEA-Quant[18, 19] was also proposed to evaluate the enrichment of a given annotation among abundant that are reproducibly quantified in a set of replicates. Such methods are capable of detecting annotations for which the members are showing a moderate level of differential expression, but not necessarily a significant one for most individual genes or proteins. Nevertheless, some of the annotations can be quite general. Indeed, GO include terms such as “cytoplasm” or “metabolism”. The relationship between the proteins that share such annotations can be quite weak. One might therefore be more interested in proteins that not only share a given annotation and show some degree of differential expression but also have some sort of relationship such as interacting together or being part of a given pathway or protein complex.

Relationships between proteins can be derived from protein-protein interactions (PPI). Indeed, proteins typically interact with one another to perform their functions, whether those interactions are transient or to form complexes. Methods such as yeast two hybrid[20], affinity purification coupled to mass spectrometry[21], and proximity labelling[22] have led to the mapping of large networks of PPIs [23–27]. Such PPI networks can therefore help to guide functional enrichment analyses by providing a structure to the data and focusing their results on annotations involving proteins with some level of relationships. We have previously developed an approach named GoNet that detected GO terms that annotate proteins that are significantly clustered in PPI networks.[28]

Herein, we propose a novel alternative approach for functional enrichment analysis: PIGNON, a Protein-protein Interaction-Guided fuNctiOnal eNrichment analysis for quantitative proteomics datasets. PIGNON investigates whether the proteins annotated by a given GO term are significantly clustered within a PPI network which is weighted by the level of differential expression of its constituting proteins. We applied this approach to a quantitative proteomics analysis of different breast cancer subtypes from Tyanova et al.[29] We compare the results of PIGNON against those of a standard functional enrichment analysis techniques and an approach performing functional enrichment analysis on proteins clusters derived from the Markov Clustering algorithm[30]. We show that our tool uniquely detects annotations whose associated proteins are significantly clustered and show a differential level of expression. Indeed, PIGNON identifies dysregulated functional annotations in the hormone receptor positive (HR+), human epidermal growth factor receptor 2 (HER2+) and triple negative (TN) breast cancer subtypes, providing insights into the molecular mechanisms of these conditions. These results demonstrate that PIGNON represents an alternative tool to standard functional enrichment analyses that can provide a more comprehensive understanding of biological samples analyzed using quantitative proteomics.

## RESULTS

We present a novel functional enrichment analysis tool named PIGNON, which identifies annotations for which the proteins are differentially expressed and clustered in a PPI network. To demonstrate PIGNON’s capabilities, we analyzed a quantitative proteomics dataset from Tyanova et al.[29], where breast cancer tumours from the HER2+, HR+ and TN subtypes were analyzed. We integrated this dataset to a PPI network for the BioGRID database[31]. The analyzed network contains 283,774 PPIs involving 16,563 proteins (see Methods for processing steps). Protein expression values are used to weight PPIs such that interacting partners with similar levels of differential expression become “closer” in the network. The clustering of 14,211 GO terms was investigated by PIGNON in the weighted PPI network. Figure 1 provides a graphical representation of PIGNON’s method.

**Figure 1:**
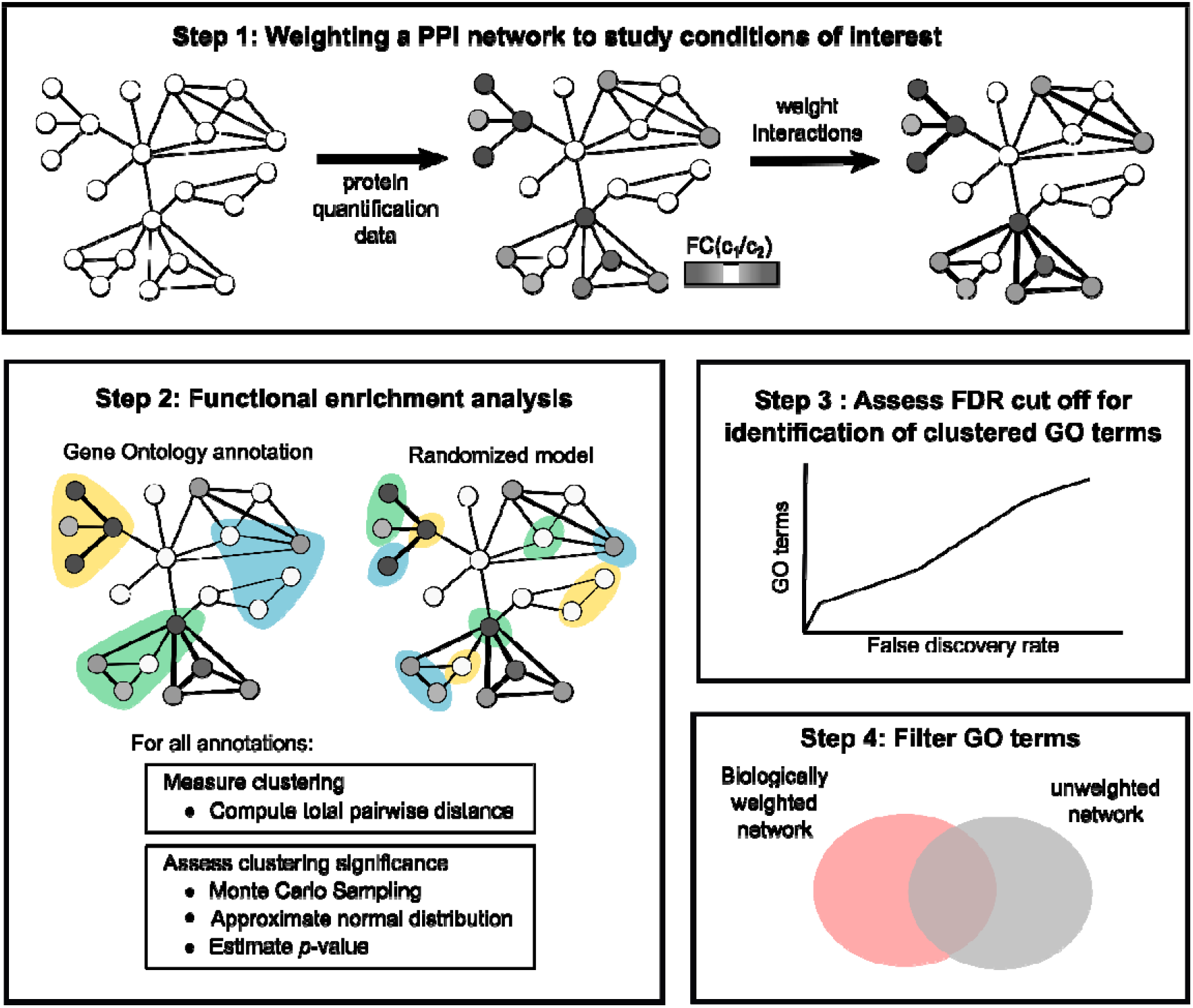
Illustrated overview of PIGNON. *Step 1.* We built the graph representation of the protein-protein interaction (PPI) network. Here, every node is a protein, and the edges between them a PPI. Edges can be weighted based on the fold change (FC) of the protein expression measured between condition 1 (c_1_) and condition 2 (c_2_). *Step 2.* We annotate the proteins of the network using Gene Ontology terms, and in parallel, the network is annotated with a shuffled set of these annotations. Here, annotations are represented by the various colours (blue, orange and green). The clustering of proteins associated to a given annotation is measured using the TPWD, a Monte Carlo approach followed by approximation of the normal distribution was performed to assess the clustering significance. *Step 3.* We assess the clustering confidence through a false discovery rate estimation. *Step 4.* Significant dysregulated GO terms are reported following filtering of significant GO terms identified in the unweighted network.

### PIGNON’s normal approximation provides accurate clustering *p*-values

Since the Monte Carlo sampling approach can hardly estimate the *p*-values of very small clustering with a p-value < 10^-7^, PIGNON estimates a null model for each protein set sizes using a normal distribution. When comparing the distribution of total pairwise distances (TPD) obtained using Monte Carlo sampling to the one generated from a normal approximation, we see that the normal approximation can be a good estimate of the Monte Carlo-derived distribution. While the fit between the two distribution is not ideal for a small protein set size such as 5, the normal distribution appears to be an upper bound on the probability of small clusterings (Additional file 1, Supplementary Figures S1A and S1B). It therefore yields a conservative estimate of the clustering *p*-values, which is desirable. The fit between the two distribution appears to increase as the protein set size grows (Additional file 1, Supplementary Figures S1C-F). Given those results, we chose to estimate clustering significance *p*-values using the normal approximation p-values.

### PIGNON can accurately estimate the statistical significance of clusterings

Given the fact that PIGNON’s *p*-values are estimations from a normal approximation, we estimate the FDR associated with a given *p*-value significance threshold. We compared the FDRs associated with the *p*-value significance threshold achieved by PIGNON using both sampling strategies (unweighted and weighted) and both types of networks (expression-weighted and unweighted). As expected, the FDRs drop as *p*-values decrease for all approaches. (Additional file 2, Supplemental Figure S2) Here, the performances of PIGNON when processing the expression-weighted and -unweighted networks are similar, which indicates that the weighting of PPIs does not greatly alter the accuracy of the p-value estimations. Interestingly, at the same *p*-value threshold, the FDRs of the weighted sampling approach are significantly lower than those of the unweighted sampling method. For instance, the weighted sampling approach yields a FDR of 0.005 at a *p*-value threshold of 0.001 when analyzing the weighted networks comparing the HER2+ subtype with TN, while the unweighted sampling method yields a FDR of 0.37 at the same *p*-value cut-off. This result is likely indicating a more accurate null model for clustering significance estimation. We observe a similar behaviour when we compare the number of significantly clustered GO terms under a given FDR threshold. (Figure 2) Specifically, the number of significant GO terms at various FDRs are comparable when PIGNON analyzes expression-weighted and unweighted networks, while the weighted sampling approach identifies more significant GO annotations than the unweighted sampling approach. For instance, the weighted sampling detected 2,212 significantly clustered GO terms in the weighted network of HER2+ subtype compared to TN at a FDR < 0.01. (Additional file 3, Supplementary Table 1) At the same FDR threshold, the unweighted sampling approach only detected 749 clustered GO terms. (Additional file 4, Supplementary Table 2) These results indicate that PIGNON can reliably identify GO terms that are significantly clustered in the PPI network. (see unweighted network results; Additional file 5, Supplementary Table 3, Additional file 6, Supplementary Table 4)

**Figure 2:**
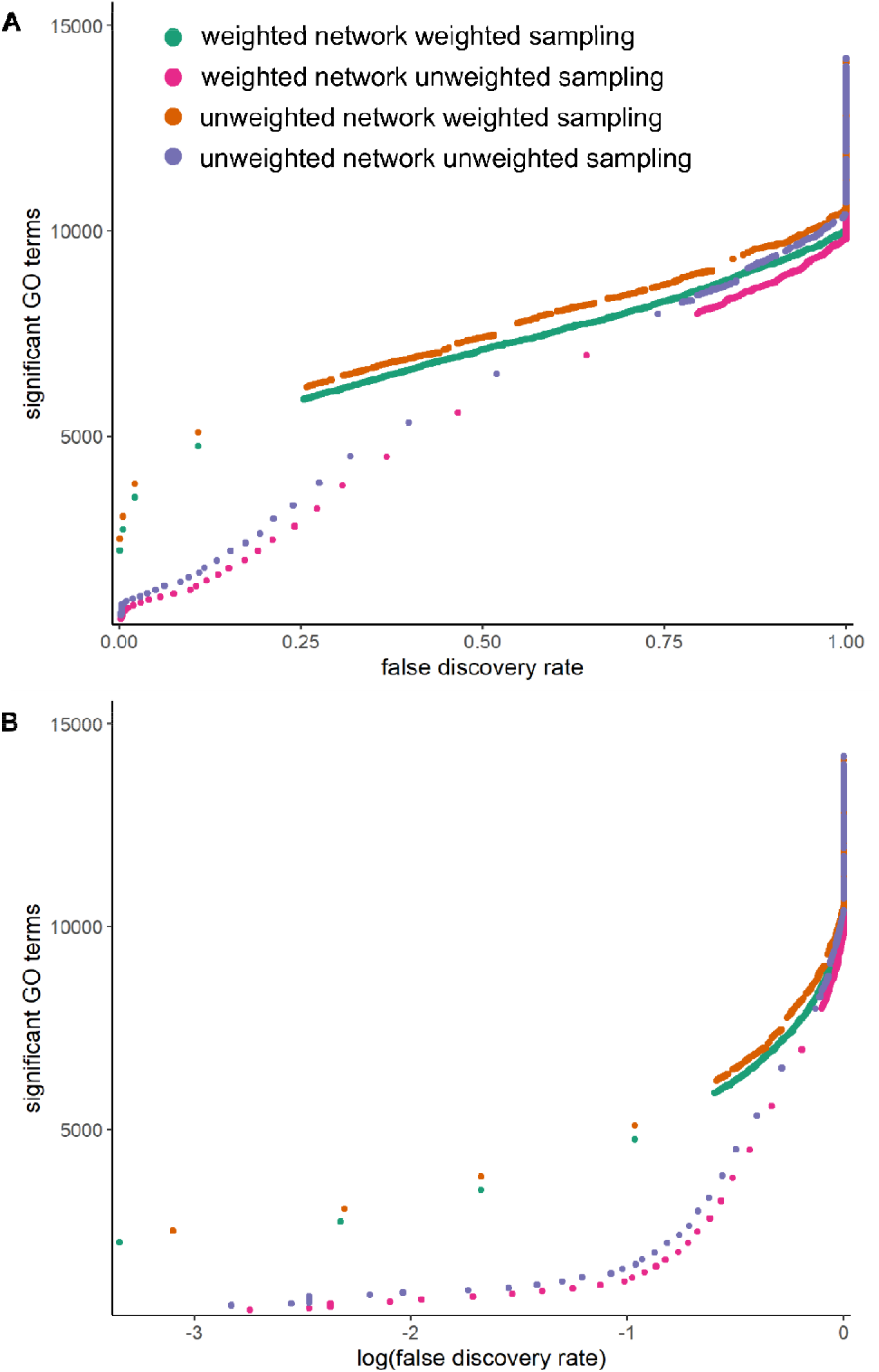
PIGNON identifies significant pathways under various conditions. Number **of** significant GO terms identified at various (A) FDR and (B) logged FDR cut-off for the vario**us** implementations of PIGNON: HER2+/TN weighted network with unweighted sampling, t**he** HER2+/TN weighted network with weighted sampling, the unweighted network w**ith** unweighted sampling, and the unweighted network with weighted sampling.

### PIGNON detects annotations for which the associated proteins are clustered in the PPI network and dysregulated

Since PIGNON evaluates for an annotation both the clustering of the associated proteins and their level of dysregulation, we looked to assess the role of the PPI network connectivity on the identification of a significant GO term. To do so, we compared the GO terms identified with our approach using the expression-weighted and unweighted PPI network for the same significance threshold. For all three breast cancer subtype comparisons, the majority of the GO terms detected were also found to be clustered in the unweighted network. Specifically, at an FDR < 0.001, 2,212 GO terms were detected in the HER2+/TN expression-weighted network, while 2,044 of these GO terms were detected to be clustered by PIGNON in the unweighted network (Figure 3A). Similar results are obtained when comparing the other breast cancer weighted networks to the unweighted network. At an FDR < 0.001, 1,934 of the 1,953 GO terms detected in the HR+/TN weighted network were found to be clustered in the unweighted network (Figure 3B) (Additional file 7, Supplementary Table 5). While 1,750 of the 1,953 GO terms detected in the HER2+/HR+ weighted network were also clustered in the unweighted network (Figure 3C) (Additional file 8, Supplementary Table 6). These shared GO terms are likely consequences of the innate clustering of their annotating proteins in the PPI network. Hence the 168 GO terms that are unique to PIGNON analysis of the HER2+/TN expression-weighted network are consequent of the dysregulation of the annotated proteins in these breast cancer subtypes. The same applies to the 19 GO terms and 203 GO that are unique to the PIGNON analysis of the HR+/TN and HR+/HER2+ expression-weighted PPI networks.

**Figure 3:**
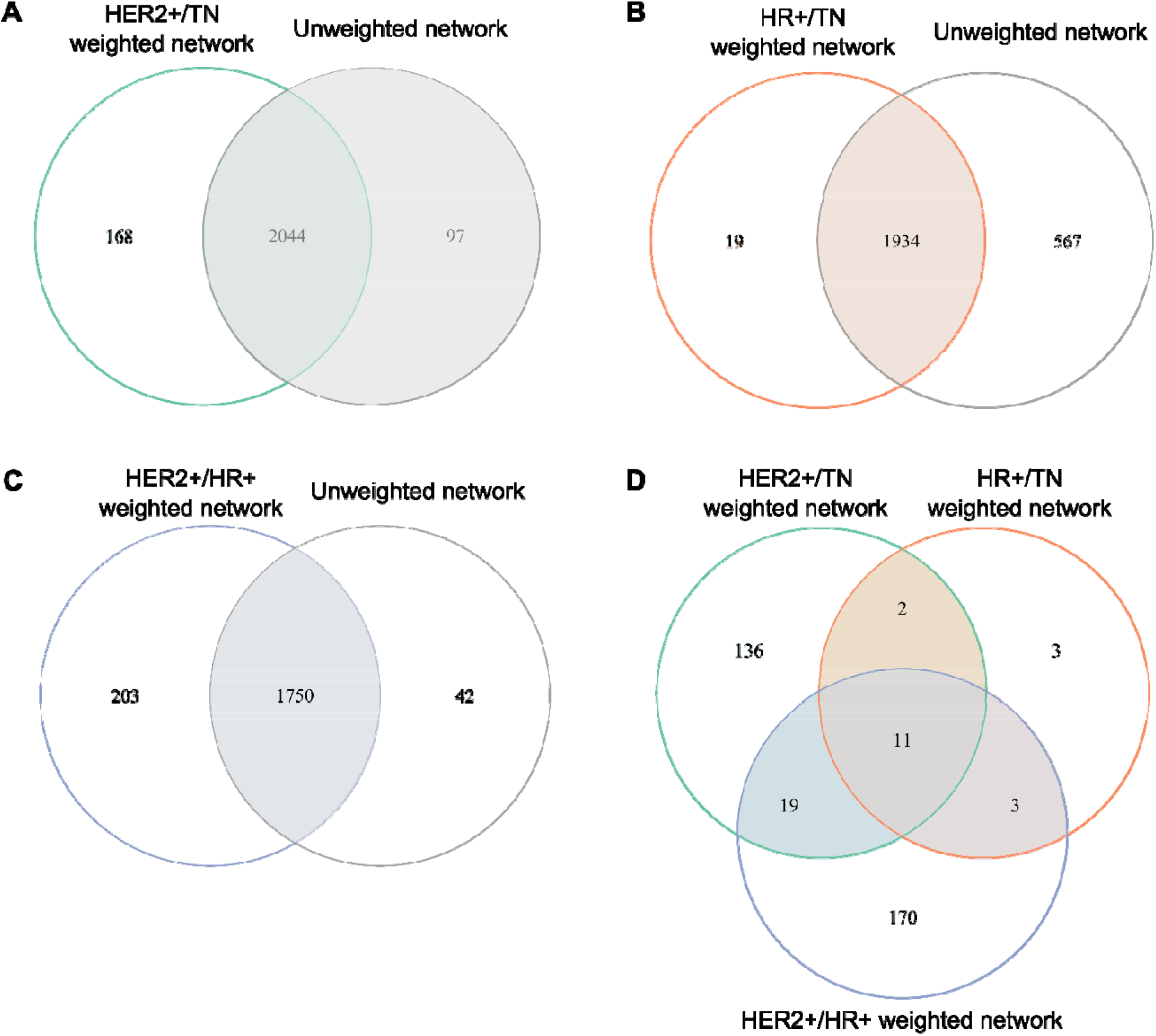
Filtering significant GO terms enables the identification of dysregulated biological processes in breast cancer subtypes. Venn diagram comparing the significant GO terms identified in the unweighted network to (A) the HER2+/TN, (B) the HER2+/HR, and (C) HR/TN weighted networks with weighted sampling. Significant GO terms are reported at an FDR < 0.001. (D) Venn diagram comparing the overlap of breast cancer subtypes.

### PIGNON detects annotations that are not highlighted by standard approaches

We first benchmarked our approach to a standard statistical pipeline, where the significance of differential protein expression is assessed, and a GO enrichment analysis is then performed on the significant proteins. This approach deemed 3 proteins to be significantly differentially expressed between HER2+ and TN as well as 4 and 0 between HER2+/HR+ and HR+/TN, respectively, at a FDR-adjusted p-value < 0.05 (Additional file 9, Supplementary File 1). From these proteins, no GO terms were found to be enriched among the differentially expressed proteins between any of the conditions (Additional file 10, Supplementary File 2).

Furthermore, we also benchmarked PIGNON against an approach combining the Markov Clustering (MCL) algorithm to identify protein clusters in the PPI network and Ontologizer to detect GO enrichments among these clusters, while considering parent-child ontologies. At comparable thresholds (i.e. FDR adjusted-p<0.001 and FDR <0.001), the MCL-Ontologizer approach identified 2,693 GO terms in the HER2+/TN expression-weighted network, while our approach identified 2,212 GO terms, of which 886 of these GO terms were identified in both approaches. (Figure 4A) (Additional file 11, Supplementary File 3) Hence, PIGNON detected 1,067 GO terms deemed to be clustered in the HER2+/TN expression weighted network that an approach using classical tools failed to detect. Similar results were obtained in the HR+/TN and HER2+/HR+ weighted network, where 2,702 and 2,722 GO terms were identified in the MCL-Ontologizer approach while 1,953 GO terms where identified in PIGNON for both networks. This translated to 893 and 971 GO terms identified in both approaches in the HR+/TN and HER2+/HR networks respectively. (Figure 4B, 4C) (Additional file 12, Supplementary File 4; Additional file 13, Supplementary file 5)These results suggest that PIGNON highlights unique information about the dataset investigated that more standard approaches do not. It further illustrates that it can be an additional approach in the data mining of quantitative datasets.

**Figure 4:**
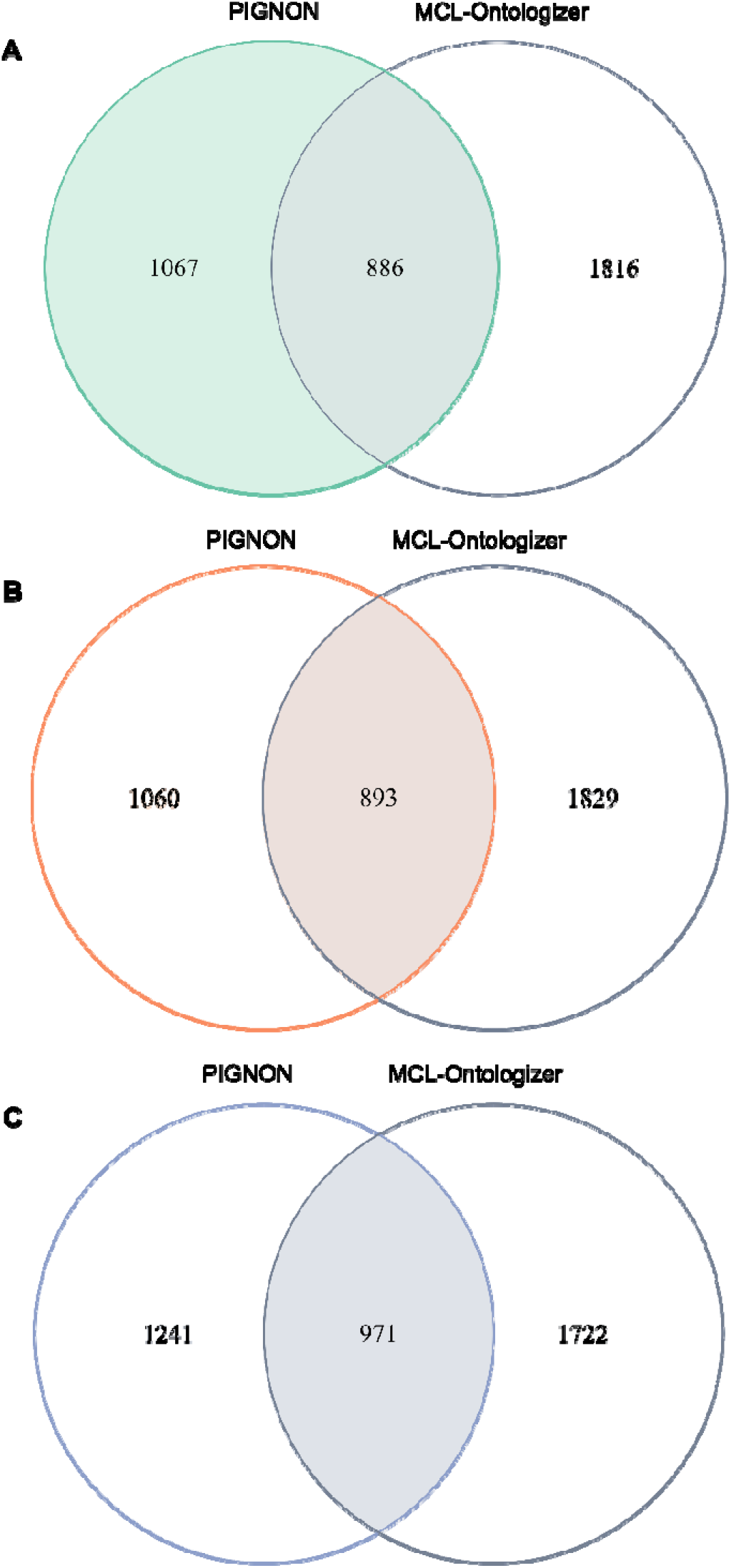
Our algorithm identifies different pathways compared to the MCL-Ontologizer approach. Venn diagram comparing the overlap of significant GO terms identified by PIGNON and the MCL-Ontologize approach on the (A) HER2+/TN, (B) HR/TN, and (C) HER2+/HR at comparable thresholds (FRD < 0.001, FDR adjusted-p < 0.001).

### PIGNON identifies GO terms that are known to be affected in different breast cancer subtypes

To limit GO terms to those of the highest biological relevance to the studied breast cancer subtypes, we filtered the identified GO terms identified by PIGNON to those that are unique to the weighted network analysis when compared to that of the unweighted network. We identified 168 breast cancer pathways dysregulated between the HER2+ and TN subtypes, 203 breast cancer pathways between HER2+ and HR+ subtypes, 19 breast cancer pathways between HR+ and TN subtypes. (Figure 3A-C) These results indicate that PIGNON identified more shared GO terms between HER2+ and HR+ subtypes, and HER2+ and TN subtype compared to the HR and TN subtypes. For the most part, the GO terms detected by PIGNON as being dysregulated have been previously characterized in breast cancers in the past but have not necessarily been associated with these particular subtypes. Pathways revealed in the HER2+ and TN subtypes include COPII vesicle coating and homotypic cell-cell adhesion (Figure 5A) whose upregulations are essential drivers of metastatic breast cancers[32, 33]. HR+ and TN were characterized by adherence junction organization whose dysregulation is implicated in lobular type breast cancers[34] and caveolin-mediated endocytosis, which is prevalent in non-HER2+ cancers [35] (Figure 5B). HER2+ and HR+ subtypes were shown to be dysregulated by Wnt signalling processes, which have been universally characterized as implicated across breast cancers. [36](Figure 5C). (For cellular components, see Additional file 14, Supplementary Figure S3; for molecular functions see Additional file 15, Supplementary Figure S4)

**Figure 5:**
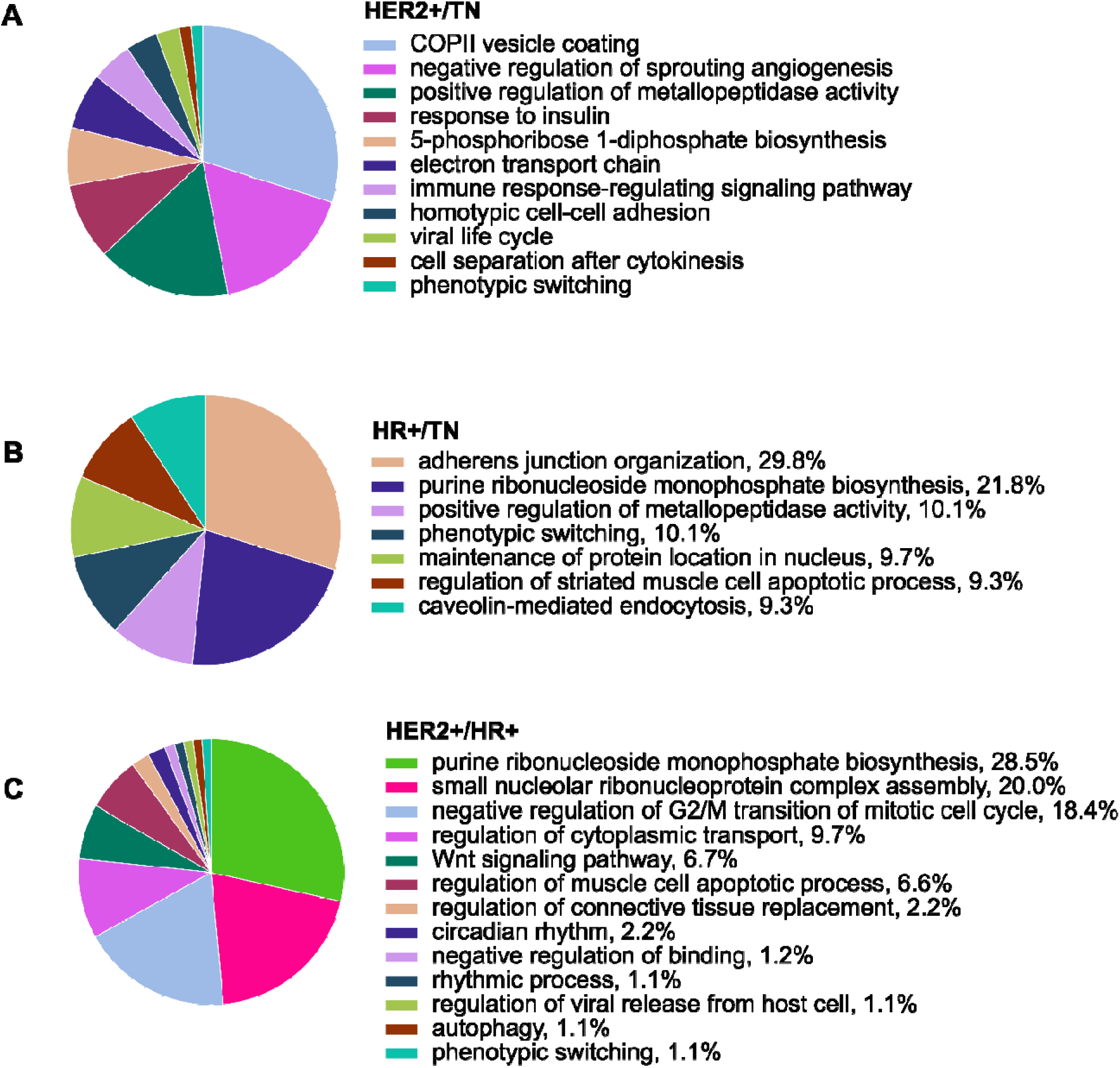
Unique significant biological processes in breast cancer subtypes. CirGO visualization of unique identified biological processes in (A) HER2+/TN, (B) HER2/HR and (C) HR/TN weighted networks at an FDR < 0.001.

We combined the results of the PIGNON analyses comparing the different breast cancer subtypes to compare the GO terms overlapping between subtypes and those that are unique to each breast cancer subtype (Figure 4D). Common pathways to all breast cancer subtypes include purine ribonucleotide triphosphate biosynthesis, the regulation of metallopeptidase activity and cortical actin cytoskeleton, which have been shown to be major regulators of breast cancer pathogenesis [37–39]. Furthermore, PIGNON highlights pathways that were previously characterized in specific breast cancer subtypes, such as cell-cell junction and plasma membrane raft which are over-expressed in TN breast cancers [40, 41]. HR+ was shown to be characterized by laminin-10 complex whose downregulation is known in HR+ breast cancer [42] and substrate adhesion dependant cell spreading, which is a major regulator of HR+ breast cancers[43]. While the activation of L-lactate dehydrogenase activity and platelet aggregation characterized in the HER2+ breast cancers are important in the metastatic processes.[44, 45] Certain pathways identified remain to the best of our knowledge unassociated with breast cancer up to now. These include the 5-phosphoribose 1-diphosphate metabolic process, which may merit further studies to investigate its potential implication in the disease.

## DISCUSSION

### Alternative PPI weighting strategies

PIGNON weights PPIs in the network based on the fold-change of their interacting proteins between the two compared conditions. PIGNON treats both low and high fold-changes the same way. This approach enables us to look at both up- and down-regulation without prioritizing a given state and focuses on the detection of annotations that are dysregulated. An alternative method could have looked at these states independently. Indeed, the PIGNON analysis could be used to detect only annotations that are down-regulated and clustered in the network or on the other hand only up-regulated. Furthermore, the fold-change is not the only measure that could be used to integrate a notion of differential expression into a PPI network. For instance, the *t-* statistic or a *p*-value resulting from a statistical test such as the Student’s *t*-test could also be used as weights in the network. Such measures also capture the variance in expression differences between the two samples.

### Alternative clustering measures

Alternative methods to the total pairwise shortest paths could have been used to provide a measure of clustering for the set of proteins annotated by a GO term. For instance, the sum of the weights of the minimum spanning trees of the proteins annotated by a GO term could be used instead. This would only consider a subset of the edges considered by the total pairwise shortest paths and would therefore be less sensitive to outliers that are very distant from the other proteins in the network. Nevertheless, the running time of this approach could become prohibitive, since minimum spanning trees need to be computed for each set of proteins being sampled. While the shortest path between all proteins is only computed once for a given network, both the shortest path-based measure and the minimum spanning tree approach consider that all proteins contribute to the differential expression and clustering of a given annotation. However, not all proteins of a given annotation need to be dysregulated for the overall annotation to be significantly dysregulated. Therefore, a clustering measure that looked only at a fraction of the most clustered proteins in an annotation, similar to the approach adopted by LESMoN[46], may allow the detection of supplemental dysregulated annotations.

### Running time considerations and possible improvements

Given the large number of samplings performed by PIGNON, its running time is considerable. Here, our Monte Carlo method sampled 10^7^ protein sets for all protein set sizes. Reducing the number of samplings by 10-fold could decrease the running time of the algorithm while still enabling the approximation of normal distribution parameters. Furthermore, we investigate GO terms that annotate as much as 1000 proteins. GO terms with such a large number of annotations are typically not of great biological interest. Reducing the maximum size of proteins sets for which PIGNON builds a null model using Monte Carlo sampling would therefore also decrease the running time of the algorithm.

### PIGNON can be applied on any quantitative datasets

PIGNON was developed to perform functional enrichments in quantitative proteomics datasets. We applied this method to a mass spectrometry investigation of breast cancer tumour subtypes; however, it can be applied on any quantitative proteomics dataset that compares a least two conditions of interest. Furthermore, while the association between quantified proteins and PPI networks is natural, PIGNON could certainly be used to perform functional enrichment analyses in quantitative transcriptomic datasets, such as those generated using RNA-sequencing. Obviously, the expression of a transcript does not necessarily equate to the translation of its corresponding protein. Nevertheless, PIGNON could still be used to identify transcripts that are differentially expressed and for which the associated proteins are clustered in PPI networks

## CONCLUSIONS

We have designed a novel PPI-guided functional enrichment analysis for quantitative proteomics datasets. Our approach, PIGNON represents an alternative strategy to identify significantly dysregulated functional annotations. Finally, PIGNON increases our ability to characterize quantitative proteomics data set and therefore helps to provide a better understanding of the processes and mechanisms at work in biological samples.

## METHODS

### Method overview

Our approach, named PIGNON, identifies differentially expressed functional annotations by assessing the clustering of proteins associated with a given GO term within an expression weighted PPI network. The network is weighted such that interacting proteins that are similarly differentially expressed are connected using edges with a small weight. Our method measures the clustering of the proteins sharing a given annotation by computing the total pairwise distance between the proteins. It then assesses the statistical significance of this clustering using a Monte Carlo sampling approach. Finally, it estimates a False Discovery Rate for the annotations deemed significantly clustered in the weighted network by PIGNON. Figure 1 provides a graphical representation of PIGNON’s method.

### Quantitative proteomics dataset

To test our methods, we used a quantitative proteomics analysis of breast tumours from three different subtypes performed by Tyanova et al. study. [29] This study assessed the proteomics profile of whole tumours diagnosed as hormone receptor positive (HR+), human estrogen receptor 2 positive (HER2+), and triple negative breast cancer (TN) subtypes using super-SILAC coupled to mass spectrometry [47]. Weights are assigned to edges based on protein differential expression between breast cancer subtypes.

### Protein-protein interaction network and expression weighted interactions

In the context of this study, we combined the above quantitative proteomics dataset with a human PPI network obtained from the BioGRID PPI public repository (version 3.4.161) [31, 48], which contains 330,754 individual PPIs and 22,733 unique proteins. Three inclusion criteria were applied to generate the complete network. Firstly, only human-to-human protein PPIs were considered, contributing to the biological validity of this study. Secondly, only PPIs that constituted the largest connected components of the BioGRID network were kept a requirement of our statistical approach. Thirdly, only proteins that interacted with less than 1000 other proteins are kept in the network, eliminating 12 overly connected proteins that condense the network and could mask clusters that are present in the network. These filters resulted in a reduced network of 16,607 proteins involved in 266,171 PPIs. This network is represented as an undirected graph, *G* — (*V,E*) where *v∈V,* the set of vertices (proteins) and *e∈E,* the set of edges connecting two vertices, (PPIs) i.e. *e — v*_1_, *v*_2_, where *v*_1_ *v*_2_ ∈ *V*.

Based on the Tyanova et al. quantitative proteomics analysis, we assign weights to the edges of the network as follows. The level of protein differential expression of a protein *v_i_*, is defined as the fold change, *f*(*v_i_*). *f*(*v_i_*) is calculated as the ratio of the average expression of *v_i_* in one experimental condition (e.g. HER2+ tumours) over the average expression of *v_i_* in another condition (e.g. TN tumours) as provided by Tyanova et al. In the event that a protein in the network *v_i_* was not quantified by Tyanova et al., *f*(*v_i_*) was set to 1, indicating no differential expression. When comparing the HER2+/TN conditions 2,883 proteins in the network have a *f*_1_(v_*i*_) ≠1, while 2,597 and 2,613 proteins were weighted in the HR+/TN and HER2+/HR+ networks respectively. Edge weights, vv(e), for all *e ∈ E* were then defined as follows:

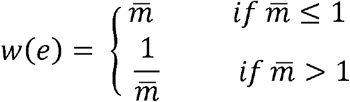

and where 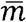 is the mean foldchange of connected vertices, v_1_ and v_2_, (i.e., the mean of *f*(*v*_1_) and *f*(*v*_2_)).

### Measuring the clustering of protein annotations in the weighted PPI network

The objective of PIGNON is to identify annotations for which the associated proteins show a level of differential expression and are clustered in a PPI network. In this study we focus on annotations derived from Gene Ontology[10]. For every GO term *T* out of the set of GO terms *O*, we measure the clustering of its associated set of proteins *S_T_* and assess their significance. Propagated GO terms, (i.e. where proteins are annotated by all of the parent GO terms of its annotations) were obtained from Himmelstein et al.(accessed 2018-10-26) [49]. PIGNON assesses the clustering of GO terms for which a minimum of 50% of their associated proteins are present in the PPI network and that annotates between 3 to 1000 proteins. This ensures a significant representation of the GO terms analyzed and also that very general GO terms, which typically have little biological interest, are not considered. The clustering of GO terms satisfying these criteria is measured using the total pairwise distance (TPD), a measured inspired by our previously published tool GoNet[28]. Formally, the TPD of the set of proteins *S_T_* associated with a GO term *T* is defined as:

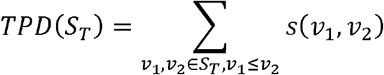

Where *s*(*v*_1_, *v*_2_) is the weighted shortest path distance between the two proteins *v*_1_ and *v*_2_. Hence proteins that show an important level of differential expression will be connected with edges with a small weight, thereby reducing their shortest path distances. The weighted shortest path between all pairs of proteins in the network using Floyd-Warshall algorithm [50, 51].

### Assessing the clustering significance of annotations in the weighted PPI network

To assess the significance of protein clustering of a GO term, PIGNON builds a null model of the TPD for each size of set of proteins *S_T_* ranging from 3 to 1000 as described above. Here we present two methods to build these null models, which are both based on a Monte Carlo sampling strategy. The first is inspired by the sampling strategy from GoNet and randomly samples proteins with a uniform probability, with the exception that only proteins annotated by at least 1 GO term can be sampled[28] and computes the TPD of each sample to build a null distribution of the TPD for a given number of proteins. We will refer to this approach as the unweighted sampling. The second is inspired by another clustering approach LESMoN[46] and samples proteins with a probability that is roughly proportional to their number of annotations. We will refer to this approach as the weighted sampling. A weighted approach enables the distribution to be more reflective of the impact of. proteins that are annotated with multiple GO term annotations can have on the clustering results, since such proteins would be influence the TPD of multiple GO terms. A null model more often selecting such proteins is therefore likely to be a more accurate model to statistical assessment. For both method 10^7^ random samplings are performed.

The main limitation of these sampling strategies is that they are unable to capture very rare events (i.e. very small TPDs with extremely small *p*-values: < 10^-7^). In order to address this issue, one can approximate the null distribution of TPDs with a distribution that has a defined cumulative distribution function. PIGNON therefore computes the TPD mean and standard deviation from the Monte Carlo samples and approximate a gaussian distribution as the null model. From the gaussian distribution, we assess the *p*-value of obtaining a given TPD α as the probability of obtaining a TPD at least as small as α when randomly selecting a set of proteins in the network.

### Estimating a False Discovery Rate for a given level of statistical significance

Given the large number of GO terms for which clustering will be assessed, multiple hypothesis testing is an important consideration. However, a large number of GO terms share many annotating proteins and therefore the clustering statistical assessment of these GO terms is far from being independent. Consequently *p*-value correction methods such as Bonferroni are likely too stringent. Instead, we restrict reported clustered annotations using an estimated False Discovery Rate (FDR).

In order to estimate this FDR, associations between GO terms and their proteins are randomized. This shuffling process takes place by selecting at random a pair of GO term-protein association and swapping the protein annotations. This process is repeated 687,855,000 times (a 1000 times the number of GO term-protein associations). In this process, GO terms that are not swapped if one of them annotates both proteins in the pair. When this process is completed the clustering statistical significance of the shuffled GO terms *O\** is assessed using the same method as above. We estimate the FDR at given *p*-value threshold, as the proportion of sets of proteins from shuffled GO terms, *S_T*_*, over the number of sets of proteins from actual GO terms, *S_T_*, with a clustering p-value smaller than *p*. Specifically, 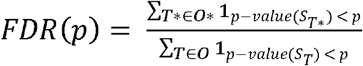.

GO terms, *S_T_*, for which the proteins are associated with a FDR < 0.01 are deemed significantly clustered in the weighted network.

### Benchmarking approaches

#### Unweighted network

PIGNON can be modified such that it solely evaluates the clustering of GO terms and does not considered any differential expression. The network effectively becomes unweighted, where all edges have a weight of 1. This unweighted network analysis is used to contrast PIGNON’s weighted network analysis.

#### Markov Clustering algorithm coupled to Ontologizer

We compare our PIGNON algorithm to a topological clustering algorithm followed by a GO enrichment analysis. We used the Markov Clustering (MCL) algorithm [30] to identify clusters in the PPI network. MCL was executed on the same network produced with the filtering procedures described above with one exception. Edge weights were inverted since MCL clusters proteins based on similarity. The inflation hyperparameter was set to 2.0. The MCL analysis yielded 1,462 clusters with greater than 3 proteins in the HER2+/TN weighted network and 1,499 clusters in both the HR+/TN and HER2+/HR+ weighted network. (Additional file 14, Supplementary File 6). These clusters were analysed using Ontologizer[14], a Gene Ontology enrichment analysis tool. Ontologizer uses a modified Fisher exact test to assess the statistical significance of the GO term enrichments among the MCL clusters using as background the 16,563 proteins in the PPI network.

When correcting for multiple hypothesis testing, we considered each GO term identified in the Ontologizer analysis in all MCL clusters as an individual statistical tests. We performed a Benjamini-Hochberg correction on the resulting *p*-values and report the smallest FDR-adjusted *p*-value of each GO term.

#### Standard functional enrichment analysis

We analyzed the Tyanova dataset using a standard statistical approach, where differentially expressed proteins are identified using a two-tail Student’s *t*-test and corrected for multiple hypothesis testing with Benjamini-Hochberg method. Ontologizer is then used to assess the level of enrichment of GO terms within the set of differentially expressed proteins, using the entire set of quantified proteins as background.

### Visualization of significant GO terms

We summarized the significant GO terms obtained by each analysis using REVIGO[52] allowing 0.7 SimRel semantic similarity to reduce the redundancy in the reported results. REVIGO’s output is then fed to the CirGO software package [53] to generate a Pie Chart visualization of the results.

## List of abbreviations

GO: Gene O ntology
PPI: protein-protein interaction
HR+: hormone receptor positive
HER2+: human estrogen receptor 2 positive
TN: triple negative breast cancer
TPD: total pairwise distance
FDR: false discovery rate
MCL: Markov Clustering

## Declarations

Ethics approval and consent to participate

Not applicable

Consent for publication

Not applicable

## Availability of data and materials

All data results generated during this study are included in this published article and its supplementary information files.

PIGNON is implemented as an open-source platform independent Java program and his available at this address: https://github.com/LavalleeAdamLab/PIGNON. The BioGRID PPI network can be download here: https://thebiogrid.org/ and the quantitative proteomics data from Tyanova et al.[29] was downloaded here: https://www.nature.com/articles/ncomms10259

## Funding

The authors acknowledge funding from the following sources: Natural Sciences and Engineering Research Council (NSERC) of Canada Discovery grants to M.L.A. R.N. held a NSERC Canada Graduate Scholarship and an Ontario Graduate Scholarship. A.B. held a Mitacs Globalink Research Award. This research was enabled by support provided by Compute Ontario (https://computeontario.ca/) and Compute Canada (www.computecanada.ca) with Resource

Allocation to M.L.A.

## Authors’ contributions

R.N. and M.L.A. designed the algorithm. R.N. and A.B. implemented the software package. R.N., A.B. and M.L.A. contributed to writing the manuscript. All authors read and approved the final manuscript.

## Acknowledgements

Not applicable

## Additional files

**Additional file 1, Supplementary Figure 1: Approximated normal distribution follow the Monte Carlo Sampling trend.** (pdf)

**Additional file 2, Supplementary Figure 2: PIGNONs FDR performance decreases as significance scores increase.** (pdf)

**Additional file 3, Supplementary Table 1: PIGNON results on HER2+/TN weighted network with weighted sampling**. (.xlsx)

**Supplemental file 4, Supplementary Table 2: PIGNON results on HER2+/TN weighted network with unweighted sampling.**

**Additional file 5, Supplementary Table 3: PIGNON results on unweighted network with unweighted sampling.** (.xlsx)

**Additional file 6, Supplementary Table 4: PIGNON results on unweighted network with weighted sampling**. (.xlsx)

**Additional file 7, Supplementary Table 5: PIGNON results on HR/TN weighted network with weighted sampling**. (xlsx)

**Additional file 8, Supplementary Table 6: PIGNON results on Her2/HR weighted network with weighted sampling.** (xlsx)

**Additional file 9, Supplementary File 1: t-test results on protein expression from Tyanova et al. dataset.** (.zip)

**Additional file 10, Supplementary File 2: Ontologizer results of significantly dysregulated proteins from t-test results in Tyanova et al. dataset.** (.zip)

**Additional file 11, Supplementary File 3: Ontologizer results on clusters identified by MCL with > 3 proteins in the HER2+/TN weighted network.** (.zip)

**Additional file 12, Supplementary File 4: Ontologizer results on clusters identified by MCL with > 3 proteins in the HR+/TN weighted network.** (.zip)

**Additional file 13, Supplementary File 5: Ontologizer results on clusters identified by MCL with > 3 proteins in the HER2+/HR+ weighted network.** (.zip)

**Additional file 14, Supplementary Figure 3: Significant cellular components in breast cancer subtypes identified by PIGNON.** (.pdf)

**Additional file 15 Supplementary Figure 4: Significant molecular functions in breast cancer subtypes identified by PIGNON.** (.pdf)

**Additional file 16, Supplementary File 6: Protein clusters identified by Markov Clustering Algorithm with >3 proteins in breast cancer weighted networks.** (.zip)

