## Supplementary Figure S1 for "PIGNON: A protein-protein interaction-guided functional enrichment analysis for quantitative proteomics"

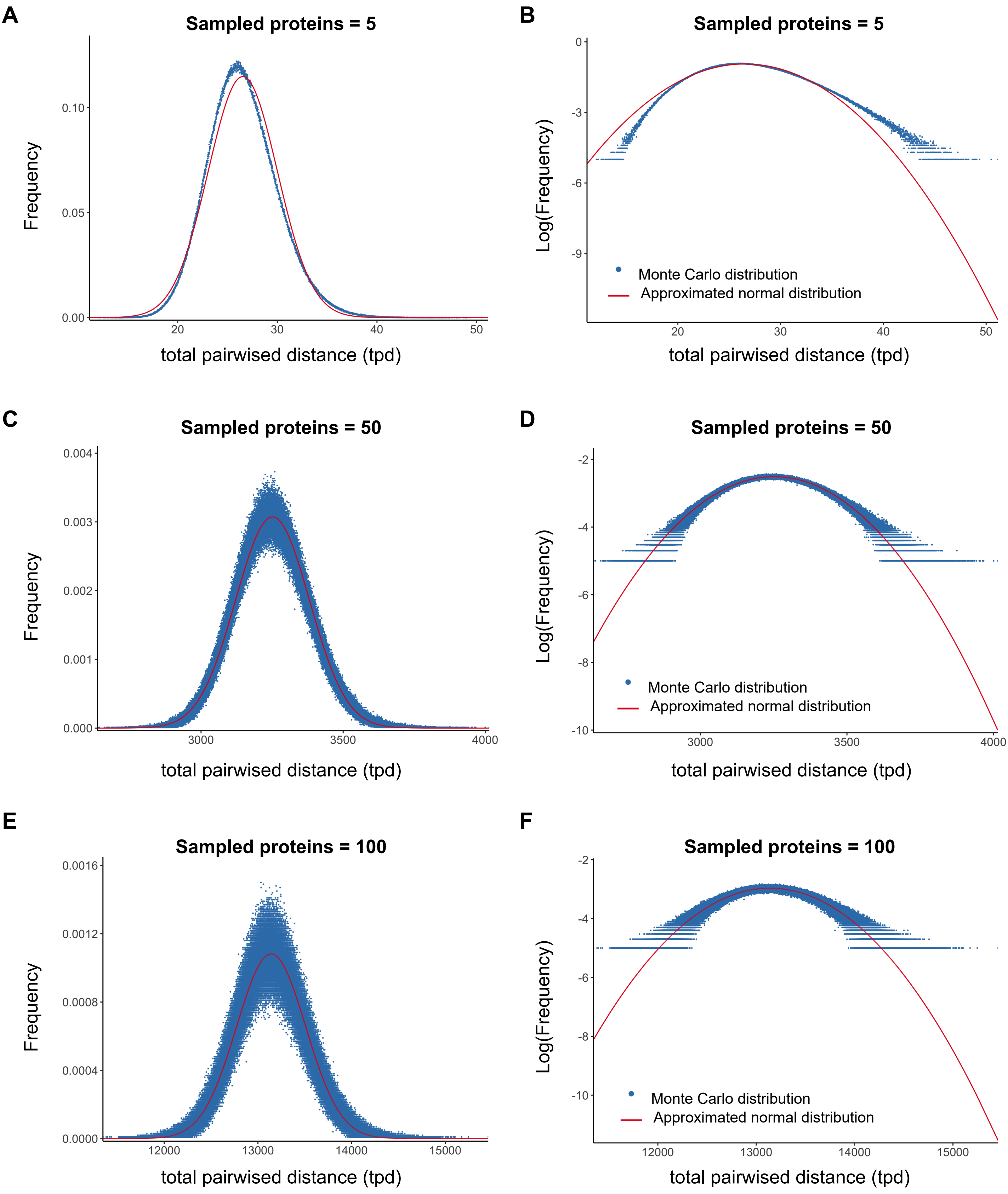


**Supplemental Figure 1: Approximated normal disributions follow the trend of Monte Carlo sampling.**Comparing the (A) frequencies and (B) logged frequencies of the Monte Carlo distributions to their approximated normal distributions for n number of proteins where n = 5, 50, 100
