## Supplementary Figure S2 for "PIGNON: A protein-protein interaction-guided functional enrichment analysis for quantitative proteomics"

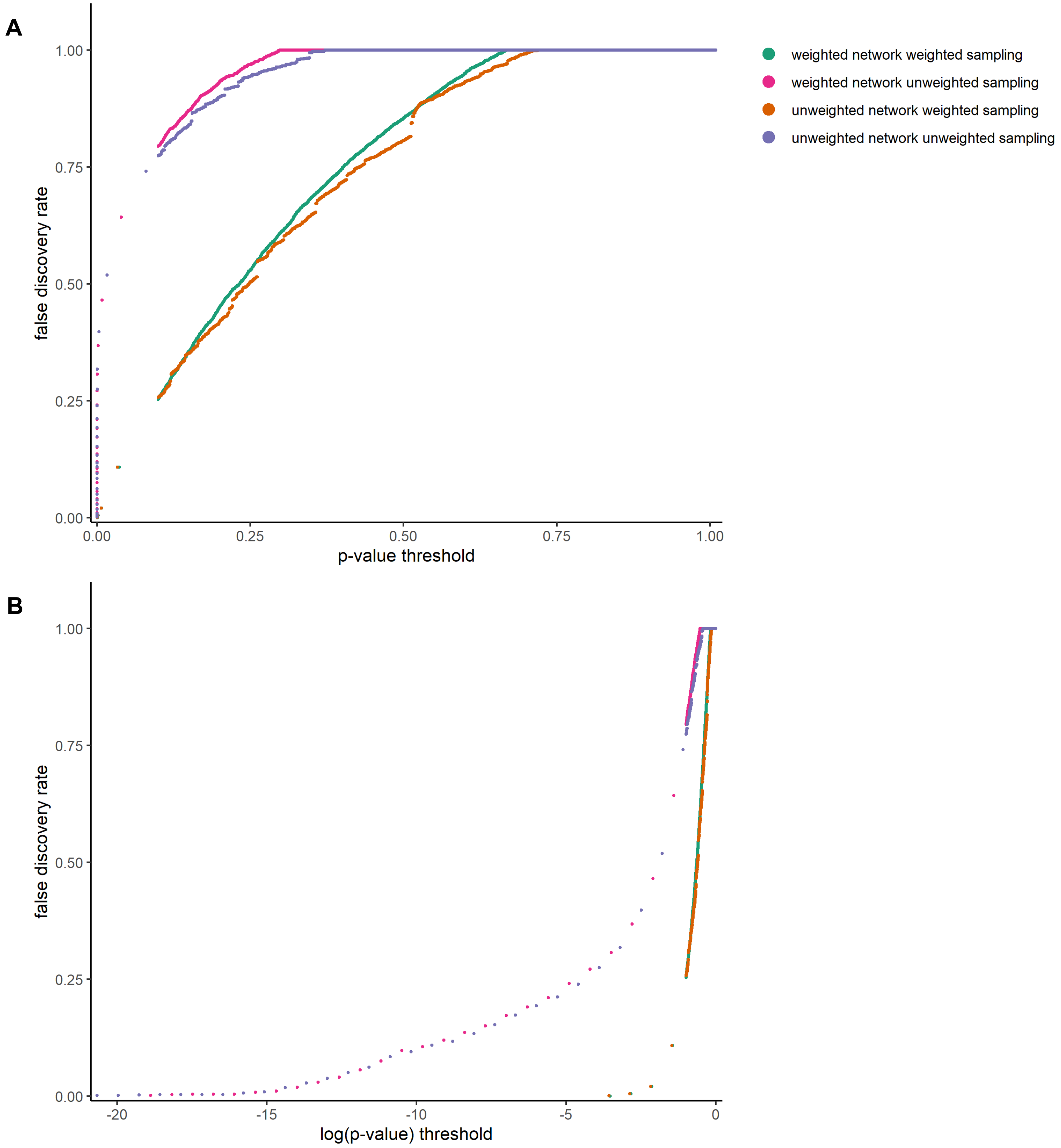


**Supplemental Figure 2: PIGNONs FDR performance decreases as significance scores increase.** FDRs calculated at various (A) p-value and (B) logged p-value thresholds for the various implementations of PIGNON: HER2+/TNBC weighted network with unweighted samp ling, the HER2+/TNBC weighted network with weighted sampling, the unweighted network with unweighted sampling, and the unweighted network with weighted sampling.
