## Supplementary Figure S3 for "PIGNON: A protein-protein interaction-guided functional enrichment analysis for quantitative proteomics"

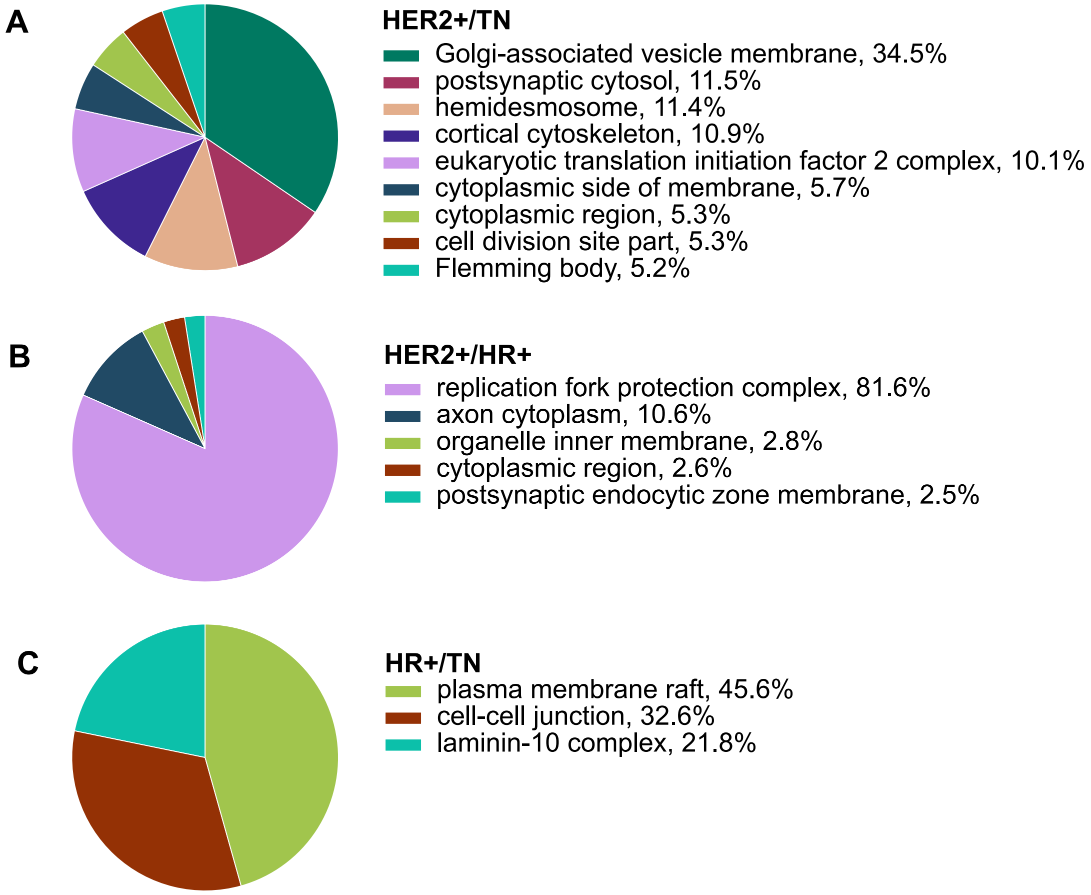


**Supplementary Figure 3: Significant cellular components in breast cancer subtypes identified by PIGNON.** CirGO visualization of unique identified cellular components in (A) HER2+/TNBC, (B) HER2+/HR and (C)HR/TNBC weighted networks at an FDR < 0.001.
