## Supplementary Figure S4 for "PIGNON: A protein-protein interaction-guided functional enrichment analysis for quantitative proteomics"

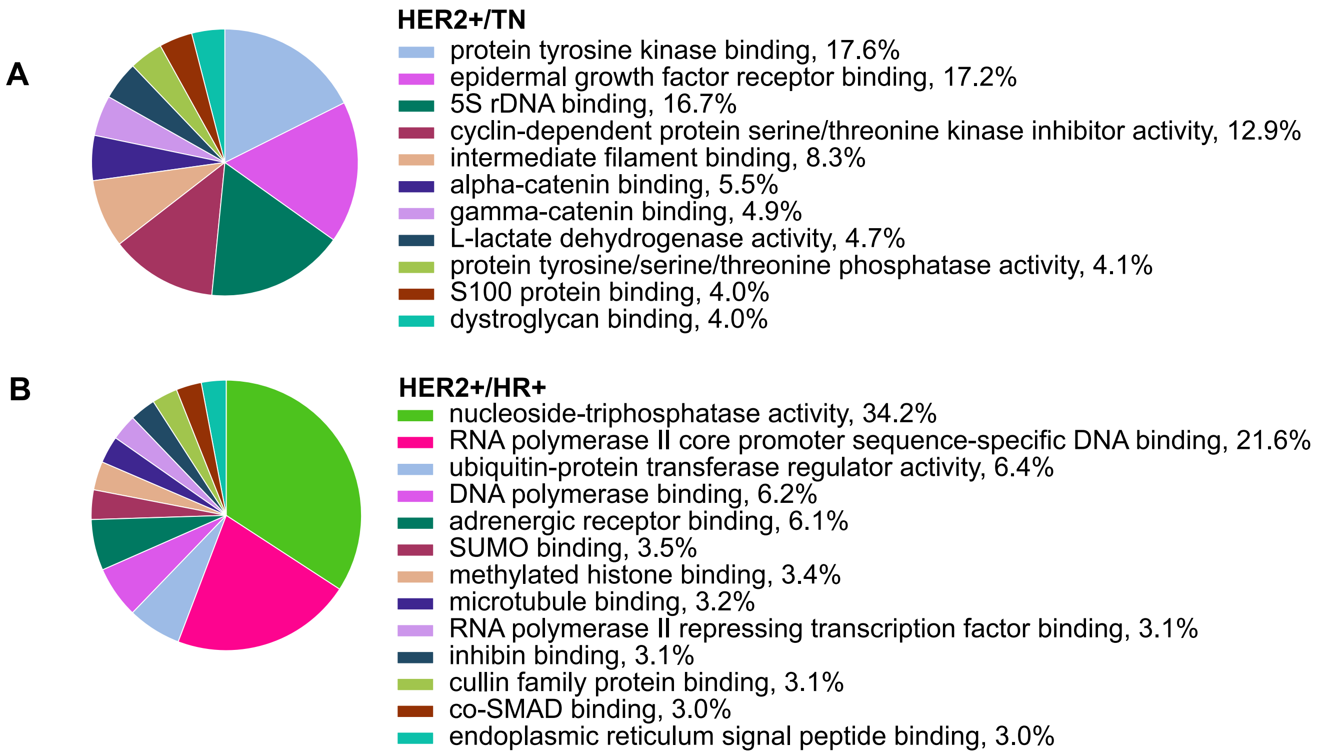


**Supplementary Figure 4: Significant molecular functions in breast cancer subtypes identified by PIGNON.** CirGO visualization of unique identified molecular functions in (A) HER2+/TNBC and (B) HER2+/HR weighted networks at an FDR < 0.001.
